# Geometry of antiparallel microtubule bundles regulates relative sliding and stalling by PRC1 and Kif4A

**DOI:** 10.1101/207696

**Authors:** Sithara S. Wijeratne, Radhika Subramanian

## Abstract

Motor and non-motor crosslinking proteins play critical roles in determining the size and stability of microtubule-based architectures. Currently, we have a limited understanding of how geometrical properties of microtubule arrays, in turn, regulate the output of crosslinking proteins. Here we investigate this problem in the context of microtubule sliding by two interacting proteins: the non-motor crosslinker PRC1 and the kinesin Kif4A. The collective activity of PRC1 and Kif4A also results in their accumulation at microtubule plus-ends (‘end-tag’). Sliding stalls when the end-tags on antiparallel microtubules collide, forming a stable overlap. Interestingly, we find that structural properties of the initial array regulate PRC1-Kif4A mediated microtubule organization. First, sliding velocity scales with initial microtubule-overlap length. Second, the width of the final overlap scales with microtubule lengths. Our analyses reveal how micron-scale geometrical features of antiparallel microtubules can regulate the activity of nanometer-sized proteins to define the structure and mechanics of microtubule-based architectures.

## Introduction

The organization of microtubules into specialized architectures is required for a diverse range of cellular processes such as cell division, growth and migration [1, 2]. Microtubule-crosslinking proteins play important roles in determining the relative orientation, size and dynamics of microtubule-based structures. These proteins include molecular motors that utilize the energy from ATP hydrolysis to mediate the transport of one microtubule over another (referred to as ‘relative sliding’) [3–5]. Motor proteins frequently act in conjunction with non-motor microtubule crosslinking proteins that oppose relative sliding and regulate both the stability and the size of the arrays [1, 2, 6]. The activities of motor and non-motor proteins are in turn modulated by the microtubule cytoskeleton. At the nanometer length-scale, numerous tubulin isotypes and post-translational modifications on tubulin act as a code to tune the activity of microtubule associated proteins (MAPs) [7–10]. In addition, it is becoming apparent that at the micron length-scale, the geometrical properties of microtubule bundles, such as orientation, filament length and overlap length, also modulate the output of motor and non-motor proteins [11–13]. Currently, we have a limited understanding of the mechanisms by which the micron-sized features of a microtubule network are ‘read’ and ‘translated’ by associated proteins.

Arrays of overlapping antiparallel microtubules form the structural backbone of diverse cellular structures. Several insights into the mechanisms underlying the assembly of such arrays have come from examining the non-motor antiparallel microtubule crosslinking proteins of the PRC1/Ase1/MAP65 family. These evolutionarily conserved proteins play an important role in organizing microtubule arrays in interphase yeast and plant cells, and subsets of spindle microtubules in dividing cells in all eukaryotes [14–19]. It is observed that these passive non-motor proteins act in concert with a number of different motor proteins, such as those of the kinesin-4, kinesin-5, kinesin-6 and kinesin-14 families [17, 18, 20–30]. A subset of these kinesins, such as Kif4A, Kif23 and Kif20, directly bind P RC1/MAP65/Ase1 family proteins [21, 24, 25, 28, 31–33]. The diversity in the properties of motor proteins that act in conjunction with the different PRC1 homologs affords a powerful model system to elucidate the biophysical principles governing the organization of antiparallel microtubule arrays. However, thus far, the mechanistic studies of PRC1-kinesin systems have mainly focused on elucidating how microtubule sliding by kinesins is regulated by PRC1 homologs [26, 34, 35]. How the initial geometry of PRC1-crosslinked microtubules modulates the activities of associated motor proteins is poorly understood.

Here we address this question by examining the relative sliding of PRC1-crosslinked antiparallel microtubules by the kinesin Kif4A. The collective activity of PRC1 and Kif4A is required for the organization of the spindle midzone, an antiparallel bundle of microtubules that is assembled between the segregating chromosomes at anaphase in dividing cells [31, 32, 36–38]. Kif4A, a microtubule plus-end directed motor protein is recruited to the midzone array through direct binding with PRC1, where it acts to suppress microtubule dynamics [25, 28, 38]. Previous *in vitro* studies with the *Xenopus Laevis* homologs of these proteins also suggest that they can drive the relative sliding of antiparallel microtubules over short distances [25]. However, microtubule sliding by Kif4A and its modulation by the geometrical features of the initial PRC1-crosslinked microtubules remains poorly characterized. In addition to sliding, it is observed that the processive movement of PRC1-Kif4A complexes and their slow dissociation from the microtubule end result in the accumulation of both proteins in micron-sized zones at the plus-ends of single microtubules (hereafter referred to as ‘end-tags’). It is observed that: (i) the movement of motor molecules is hindered at end-tags formed on single microtubules, likely due to molecular crowding and (ii) the size of end-tags increases with microtubule length [28]. How the length-dependent accumulation of PRC1-Kif4A molecules on single microtubules impacts the organization of antiparallel bundles is unknown.

Here we show using TIRF-microscopy based assays that the collective activity of PRC1 and Kif4A results in relative microtubule sliding and concurrent end-tag formation on antiparallel microtubules. Interestingly, we find that PRC1-Kif4A end-tags act as roadblocks to prevent the complete separation of sliding microtubules. Consequently, sliding and stalling of antiparallel microtubules by PRC1 and Kif4A result in the assembly of a stable overlap that is spatially restricted to the filament plus-ends. Surprisingly, quantitative examination of the data reveals that two aspects of the PRC1-Kif4A mediated microtubule organization are modulated by the initial geometry of crosslinked microtubules. First, the sliding velocity in this system scales with the initial length of the antiparallel overlap. Second, the size of the final stable antiparallel overlap established by PRC1 and Kif4A scales with the lengths of the crosslinked microtubules. Together with computational modeling, our observations provide insights into the principles by which the geometrical features of antiparallel arrays can be translated to graded mechanical and structural outputs by microtubule associated motor and nonmotor proteins.

## Results

### Collision of PRC1-Kif4A end-tags on sliding microtubules results in the formation of antiparallel overlaps of constant steady-state length

To investigate microtubule sliding in the PRC1-Kif4A system, we reconstituted the activity of the kinesin Kif4A on a pair of antiparallel microtubules crosslinked by the non-motor protein PRC1. For these studies, we adapted a Total Internal Reflection Fluorescence (TIRF) microscopy-based assay that we have previously used to examine relative sliding of PRC1-crosslinked microtubules by the motor-protein Eg5 [34]. First, biotinylated taxol-stabilized microtubules, labeled with rhodamine, were immobilized on a glass coverslip. Next, unlabeled PRC1 (0.2 nM) was added to the flow chamber and allowed to bind the immobilized microtubules. Finally, rhodamine-labeled non-biotinylated microtubules were flowed into the chamber to generate microtubule ‘sandwiches’ crosslinked by PRC1 on the glass coverslip (Fig. 1A). After washing out the unbound proteins, the final assay buffer containing Kif4A-GFP, PRC1 and ATP at specified concentrations was flowed into the chamber to initiate end-tag formation and microtubule sliding (Fig. 1A). Near-simultaneous multi-wavelength imaging of rhodamine-labeled microtubules and Kif4A-GFP showed that Kif4A preferentially accumulates in the overlap region of PRC1-crosslinked microtubules (Figs. 1B-D; *t* = 0 s; 0.2 nM PRC1 + 6 nM Kif4A-GFP). This is in agreement with prior findings that PRC1 selectively accumulates at regions of antiparallel microtubule overlap regions and recruits Kif4A to these sites [25, 34]. In the example shown in Figs. 1B-D, the average fluorescence intensity of Kif4A-GFP in the microtubule overlap region is 5-fold higher than the fluorescence intensity in the non-overlapped region at the first time point recorded (Figs. 1B-E; *t* = 0 s). In addition, time-lapse imaging shows an enhanced accumulation of Kif4A-GFP at the plus-ends of both the crosslinked microtubules. We refer to this region of high protein density at microtubule plus-ends as ‘end-tags’ (Figs. 1B-E; *t* = 10–40 s; ~2.5 fold enrichment of Kif4A-GFP at end-tags over the untagged overlap at 10 s). These data indicate that under these conditions Kif4A-GFP-containing end-tags are established at the plus-ends of crosslinked microtubules.

**Fig. 1.**
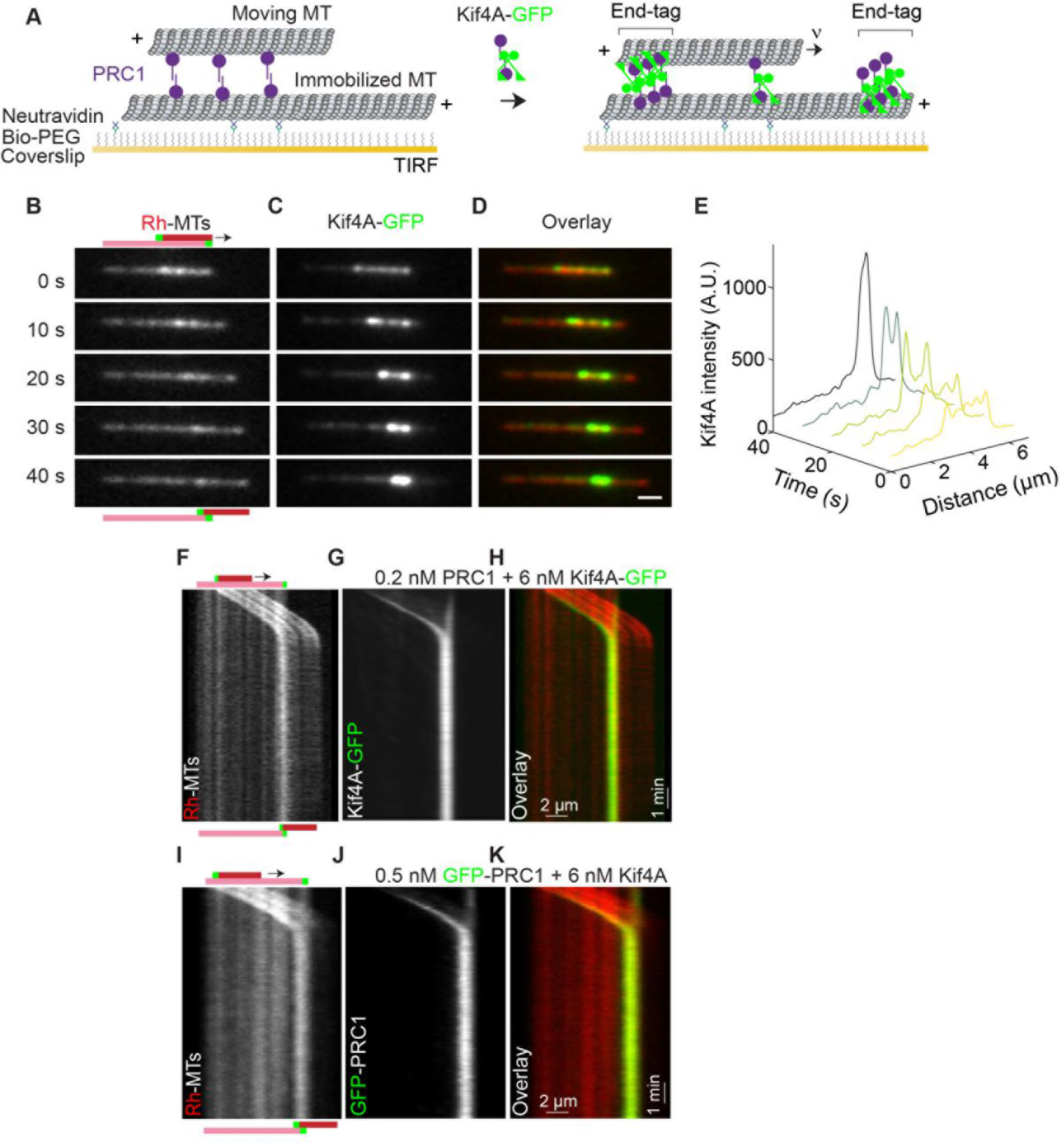
Relative microtubule sliding and the formation of stable antiparallel microtubule overlaps by PRC1 and Kif4A. **(A)** Schematic of the *in vitro* assay. A biotinylated microtubule (‘immobilized MT’, X-rhodamine labeled) immobilized on a PEG coated coverslip and a non-biotinylated microtubule (‘moving MT’, X-rhodamine-labeled) are crosslinked in an antiparallel orientation by PRC1 (purple). Microtubule sliding and end-tag formation are initiated by addition of Kif4A-GFP (green), PRC1 and ATP. **(B-D)** Representative time-lapse fluorescence micrographs of relative microtubule sliding in experiments with 0.2 nM PRC1 and 6 nM Kif4A-GFP. Images show (B) a pair of X-rhodamine-labeled microtubules, (C) Kif4A-GFP, and (D) overlay images (red, microtubules; green, Kif4A-GFP). The schematic in (B) illustrates the position and relative orientation of both the immobilized (pink) and moving (red) microtubules and the end-tags (green) at the beginning and end of the time sequence. Scale bar: *x*: 2 μm and *y*: 1 min. **(E)** Line scan analysis of the Kif4A intensity from the micrographs in (C) show the distribution of Kif4A within the overlap at the indicated time points. **(F-H)** Kymographs show the relative sliding and stalling of antiparallel microtubules (F), associated Kif4A-GFP (G) and the overlay image (red, microtubules; green, Kif4A-GFP) (H). Assay condition: 0.2 nM PRC1 and 6 nM Kif4A-GFP. Scale bar: *x*: 2 μm and *y*: 1 min. **(I-K)** Kymographs show the relative sliding and stalling of antiparallel microtubules (I), associated GFP-PRC1 (J) and the overlay image (red, microtubules; green, GFP-PRC1) (K). Assay condition: 0.5 nM GFP-PRC1 and 6 nM Kif4A. Scale bar: *x*: 2 μm and *y*: 1 min.

Time-lapse imaging and kymography-based analyses revealed that the end-tagged antiparallel microtubules slide relative to each other (Figs. 1B-D and 1F-H). Strikingly, we find that microtubule sliding stalls when the end-tags arrive at close proximity (Figs. 1B-D and 1F-H). This results in the formation of stable antiparallel overlaps that maintain a constant steady-state width for the entire duration of the experiment (Figs. 1B-D and 1F-H; *t* = 10 mins). Under these experimental conditions, we do not observe any event where the moving microtubule slides past the end-tag of the immobilized microtubule. We rarely (5%) observe sliding microtubules stall before they arrive at the plus-end of the immobilized microtubule. These observations indicate that the formation of stable antiparallel overlaps is due to the end-tags on the crosslinked microtubule pair arriving at close proximity during relative sliding.

We next examined PRC1 localization on sliding microtubules by conducting experiments similar to that described above, except with GFP-labeled PRC1 and unlabeled Kif4A (Figs. 1I-K; 0.5 nM GFP-PRC1 + 6 nM Kif4A). We find that the localization pattern of GFP-PRC1 is similar to Kif4A with the highest fluorescence intensity at the end-tags, intermediate intensity at the untagged microtubule overlap regions and the lowest intensity on single microtubules. Similar to the observations in Figs. 1F-H, we find that sliding microtubules stall when their end-tags arrive in close proximity (Figs. 1I-K).

Together, these observations suggest that human PRC1-Kif4A complexes can drive the relative sliding of antiparallel microtubules over the distance of several microns. However, sliding comes to a halt at microtubule plus-ends resulting in the formation of stable antiparallel overlaps of constant steady state length.

### Characterization of relative microtubule sliding in the PRC1-Kif4A system

To further characterize relative sliding in mixtures of PRC1 and Kif4A, we quantitatively examined the microtubule movement observed in these experiments. Analysis of the instantaneous velocity during microtubule sliding (Figs. 2A-C; 0.2 nM PRC1 + 6 nM Kif4A-GFP), reveals three phases: (1) initial sliding at constant velocity, (2) reduction in sliding velocity as the end-tags arrive at close proximity, and (3) microtubule stalling and the formation of stable overlaps that persist for the duration of the experiment.

We first focused on microtubule movement in phase-1 and investigated how the relative solution concentrations of the motor and the non-motor protein impact the initial sliding velocity. This is particularly interesting in the case of PRC1-Kif4A system as the recruitment of Kif4A to microtubules is dependent on PRC1 [25, 28]. Therefore, one possible outcome is that motor-protein movement is sterically hindered at higher PRC1 concentrations resulting in lower sliding velocities. Alternatively, it is possible that more Kif4A is recruited to microtubule overlaps at higher PRC1 concentrations and this could counter the potentially inhibitory effects of PRC1. To distinguish between these mechanisms, we compared the maximum microtubule sliding velocity (computed as the average velocity from phase-1) at two different PRC1:Kif4A concentration ratios (Fig. 2D). We found that increasing the PRC1 solution concentration 5-fold (0.2 and 1 nM) at constant Kif4A-GFP concentration (6 nM) resulted in a 4-fold reduction in the microtubule sliding velocity (velocity = 60 ± 17 nm/s at 0.2 nM and velocity=15 ± 8 nm/s at 1 nM). Similarly, in assays with GFP-PRC1 and unlabeled Kif4A, we found that increasing PRC1 concentration 2-fold (0.5 and 1 nM) at constant Kif4A levels (6 nM) resulted in a ~2-fold reduction in the microtubule sliding velocity (Supp. Fig. 1A; velocity=35 ± 13 nm/s at 0.5 nM and velocity=18 ± 6 nm/s at 1 nM PRC1). Interestingly, in these experiments, we could restore the sliding velocity by compensating the 2-fold increase in the PRC1 concentration with a 2-fold increase in the Kif4A concentration (Supp. Fig. 1A).

**Fig. 2.**
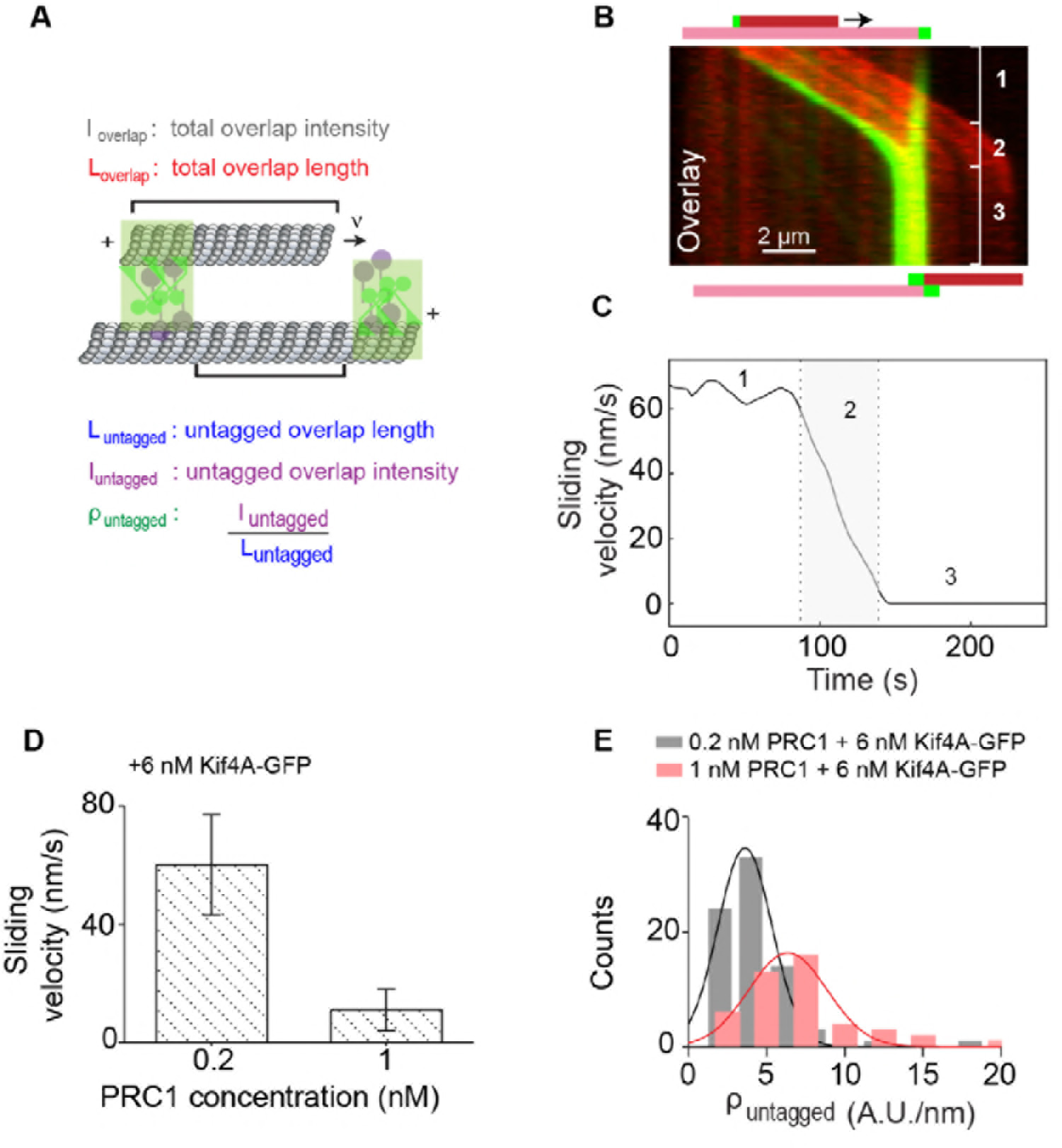
Quantitative analysis of microtubule sliding in the PRC1-Kif4A system. **(A)** Schematic of a pair of crosslinked microtubules showing the parameters described in Figs. 2 and 3. **(B)** Kymograph shows the relative sliding and stalling in a pair of antiparallel microtubules (red) and associated Kif4A-GFP (green). Assay condition: 0.2 nM PRC1 and 6 nM Kif4A-GFP. Scale bar: 2 μm. The schematic illustrates the position and relative orientation of both the immobilized (pink) and moving (red) microtubules and the end-tags (green) at the beginning and end of the time sequence. **(C)** Time record of the instantaneous sliding velocity of the moving microtubule derived from the kymograph in (B). The dashed lines demarcate the three phases observed in the sliding velocity profile: (1) constant phase, (2) slow down and (3) stalling. **(D)** Bar graph of the average sliding velocity calculated from the constant velocity movement in phase-1. Assay conditions: (i) 0.2 nM and 6 nM Kif4A-GFP (mean: 60 ± 17; N=98) (ii) 1 nM PRC1 and 6 nM Kif4A-GFP (mean: 15 ± 8; N=45). Error bars represent the standard deviation of the data. **(E)** Histograms of the initial GFP-fluorescence density in the untagged region of the overlap, *ρ*_*untagged*_. Assay conditions: (i) 0.2 nM PRC1 and 6 nM Kif4A-GFP (black; mean: 3.7 ± 1.7 A.U./nm; N=64) and (ii) 1 nM PRC1 and 6 nM Kif4A-GFP (red; mean: 6.5 ± 1.9 A.U./nm; N=33). The mean and error values were obtained by fitting the histograms to a Gaussian distribution.

One possible explanation for the reduced velocity at the higher PRC1:Kif4A concentration is that there are fewer Kif4A molecules in the overlap due to competition from PRC1 for binding sites on the microtubule surface. Therefore, we compared the Kif4A-GFP density in the untagged overlap at two different solution PRC1 concentrations (Fig. 2E). The data show that a 5-fold increase in the PRC1 concentrations results in a 2-fold increase in the average Kif4A density in the untagged overlap region, indicating that Kif4A is effectively recruited to antiparallel overlaps at the highest PRC1 concentrations in our assays (Fig. 2E). Together, these results are consistent with a mechanism in which the solution concentration of PRC1:Kif4A sets the sliding velocity by determining the relative ratio of sliding-competent PRC1-Kif4A complexes to sliding-inhibiting PRC1 molecules in the antiparallel overlap.

### Microtubule sliding velocity in the PRC1-Kif4A system scales with initial overlap length

We next examined if the initial width of the PRC1-crosslinked anti-parallel overlap impacts the sliding velocity. Remarkably, analysis of three different datasets suggests that antiparallel microtubules with longer initial overlaps slide at a higher velocity than microtubules with shorter initial overlaps under the same experimental condition (Figs. 3A-B). Note: no obvious trend was observed at the higher PRC1 concentration, possibly due to the high scatter in the data and the low sliding velocities (Fig. 3A; red squares). We also analyzed the data to determine if the microtubule sliding velocity depended on the amount of Kif4A-GFP at end-tags. However, no clear correlation was observed between these parameters using the same dataset where the sliding velocity depends on the initial overlap length (Supp. Fig. 1B; 0.2 nM PRC1 + 6 nM Kif4A-GFP, grey circles). These data suggest that the activity of PRC-Kif4A molecules in the untagged region of the antiparallel microtubule overlap is likely responsible for overlap length-dependent sliding velocity.

**Fig. 3.**
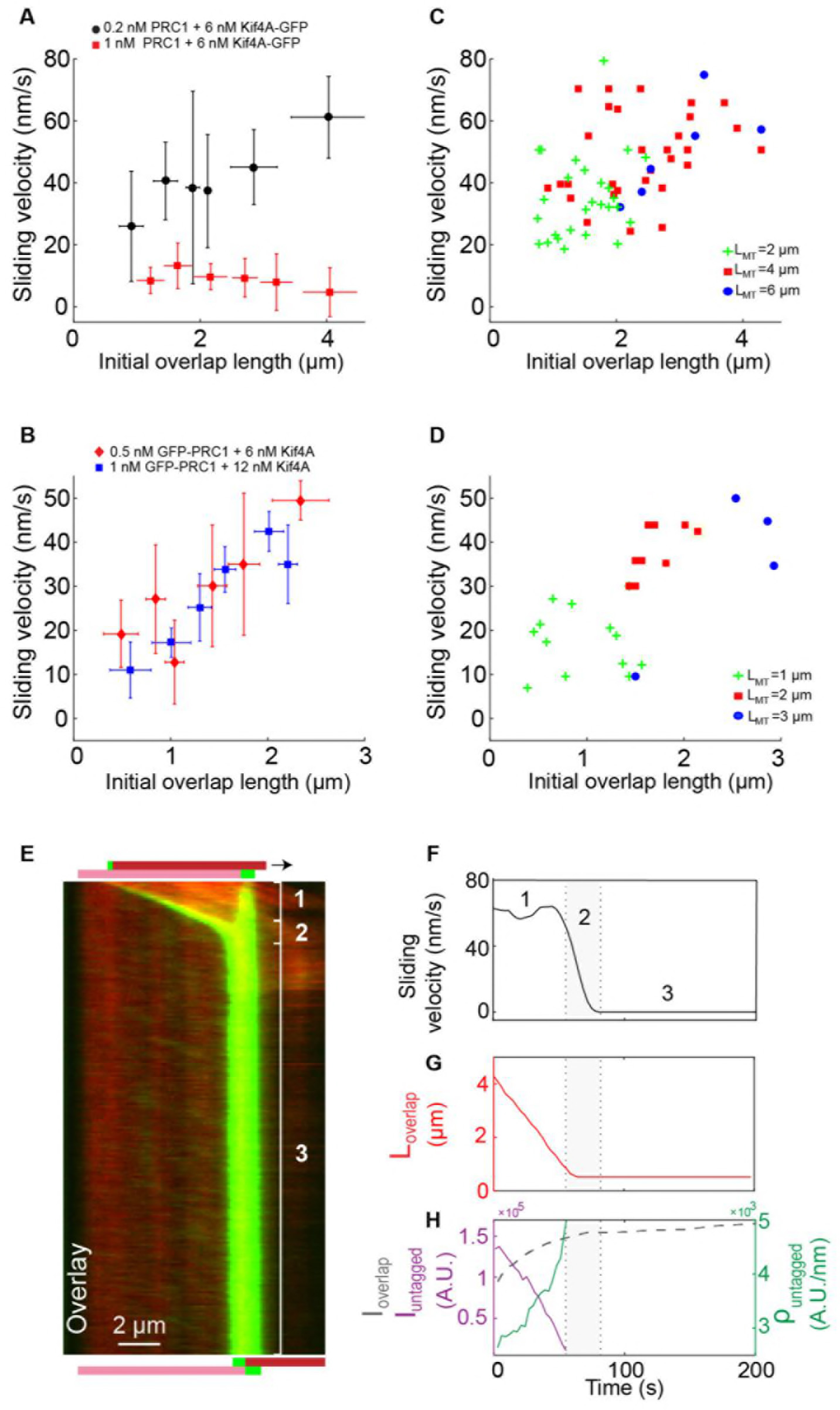
Microtubule sliding velocity in the PRC1-Kif4A system scales with initial overlap width. **(A-B)** Binned plots of initial sliding velocity versus the initial overlap length. The initial overlap length between the moving MT and immobilized MT is calculated from the rhodamine MT channel. Sliding velocity is calculated from the constant velocity movement in phase-1. (A) Assay conditions: (i) 0.2 nM PRC1 and 6 nM Kif4A-GFP (black; N=60; Pearson’s correlation coefficient=0.54) and (i) 1 nM PRC1 and 6 nM Kif4A-GFP (red; N=42; Pearson’s correlation coefficient=0.03). (B) Assay conditions: (i) 0. 5 nM GFP-PRC1 and 6 nM Kif4A (red; N=25; Pearson’s correlation coefficient=0.69) and (ii) 1 nM GFP-PRC1 and 12 nM Kif4A (blue; N=20; Pearson’s correlation coefficient=0.74). **(C-D)** Scatter plot of the average sliding velocity versus the initial overlap length color-coded by moving microtubule length, *L*_*MT*_. (C) Assay condition: 0.2 nM PRC1 and 6 nM Kif4A-GFP (green: *L*_*MT*_ = 2 ± 0.5 μM, red: *L*_*MT*_ = 4 ± 0.5 μM, blue: *L*_*MT*_ = 6 ± 0.5 μM; N=60). (D) Assay condition: 0.5 nM GFP-PRC1 and 6 nM Kif4A (green: *L*_*MT*_ = 1 ± 0.5 μM, red:*L*_*MT*_ = 2 ± 0.5 μM, blue: *L*_*MT*_ = 3 ± 0.5 μM; N=25). **(E)** Kymograph shows the relative sliding and stalling in a pair of antiparallel microtubules (red) and associated Kif4A-GFP (green). Assay condition: 0.2 nM PRC1 and 6 nM Kif4A-GFP. Scale bar: 2 μm. The schematic illustrates the position and relative orientation of both the immobilized (pink) and moving (red) microtubules and the end-tags (green) at the beginning and end of the time sequence. **(F)** Time record of the instantaneous sliding velocity of the moving microtubule derived from the kymograph in (E). The dashed lines demarcate the three phases observed in the sliding velocity profile: (1) constant phase, (2) slow down and (3) stalling. **(G)** Time record of the overlap length (red; *L*_*overlap*_) derived from the kymograph in (E). **(H)** Time record of the total fluorescence intensity in the antiparallel overlap (dashed gray; *I*_*overlap*_), fluorescence intensity in the untagged region of the overlap (solid purple; *I*_*untagged*_, and fluorescence density (intensity per unit overlap length) in the untagged region of the overlap (solid green; *ρ*_*untagged*_) derived from the kymograph in (E).

In order to separate the effect of microtubule length versus antiparallel overlap length on the sliding velocity, we re-plotted the data in Figs. 3A-B based on the length of the moving microtubule (i.e. the filament subjected to viscous drag). A scatter plot of the sliding velocity as a function of initial overlap length color-coded by the moving-microtubule length shows that longer microtubules typically form longer initial overlaps that exhibit faster sliding (Figs. 3C-D). However, the observation that long microtubules that form short overlaps exhibit slower sliding than long microtubules that form long overlaps (for example: 6 μm microtubules with ~3 μm overlap in Fig. 3C, blue dots), suggests the dominant contribution to the sliding velocity is from the initial overlap length (Figs. 3C-D). Our findings indicate that the initial length of the antiparallel overlap can tune the microtubule sliding velocity, such that longer overlaps slide at a faster rate than shorter microtubule overlaps.

The finding that sliding velocity scales with initial overlap length raised another question: is the abrupt shift from constant to decreasing sliding velocity (phase-1 to phase-2) observed in these experiments due to the transition from constant to decreasing overlap length as the moving microtubule slides past the immobilized microtubule? Such a mechanism has been described previously for the Ase1 and Ncd system (*S. pombe* PRC1 and Kinesin-14 homologs) [26]. To answer this question, we compared the time-dependent changes in the sliding velocity with total overlap length (*L*_*overlap*_) found no obvious correlation between these parameters. For example, in the kymograph shown in Fig. 3E, the reduction in microtubule overlap length begins at 0s (Fig. 3G, solid red line) but a significant reduction in the velocity is not seen until 60s (Fig. 3F, solid black line; see also Supp. Figs. 1J-L). Instead, the data suggest that the transition from sliding to stalling coincides with the end-tags on the moving and immobilized microtubules arriving at close proximity (Figs. 3E-F).

The observation that microtubule sliding occurs at a constant velocity even as the overlap shrinks, raises the following question: do the number of Kif4A molecules in the overlap change during relative sliding? Analyses of GFP intensity versus time showed that in phase-1, the total amount of Kif4A-GFP in the microtubule overlap (*I*_*overlap*_ = *I*_*end-tagged*_ + *I*_*untagged*_) initially increases and then reaches a constant level that is maintained during all three phases (dashed gray line) (Fig. 3H and Supp. Fig. 1L). This result suggests that Kif4A is retained in the shrinking overlap during sliding. Is this retention entirely due to end-tag formation or is there retention of Kif4A molecules in the untagged overlap during microtubule sliding? To answer this question, we quantitatively analyzed the Kif4A-GFP levels in the untagged region of the overlap during sliding. We find that while the total Kif4A-GFP levels in the untagged overlap region (*I*_*untagged*_) (Fig. 3H; solid purple curve) decrease with shrinking overlap, the Kif4A-GFP density (*ρ*_*untagged*_) (Fig. 3H; solid green curve; fluorescence intensity/pixel) increases two-fold with time. These findings suggest that as the overlap shrinks due to sliding, a fraction of the motor molecules is retained in the untagged region of the microtubule overlap, and possibly contributes to maintaining a constant sliding velocity.

Together, these data indicate that the velocity of microtubule sliding in the PRC1-Kif4A system is determined by the initial width of the PRC1-crosslinked antiparallel overlap. The microtubule-movement can subsequently proceed at a constant velocity, even as the overlap shrinks, possibly through the concentration of motor molecules within the overlap during relative sliding.

### The size of stable antiparallel overlaps established by PRC1 and Kif4A are determined by microtubule length and protein concentration

Our findings suggest that in the PRC1-Kif4A system, a stable antiparallel overlap is formed when the end-tags on both microtubules merge during relative sliding. We hypothesized that if stable overlaps form upon the collision of end-tags formed on the moving and immobilized microtubules, then the final overlap length (*L*_*FO*_) should be determined by the sum of the two end-tag lengths (*L*_*ET1*_ + *L*_*ET2*_) (Fig. 4A). Consistent with this, the average ratio of the 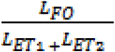 at 1 nM PRC1 + 6 nM Kif4A-GFP is ~1 (Fig. 4B, red). Similar results were observed when the experiments were performed under three different conditions with GFP-PRC1 and untagged Kif4A (Supp. Fig. 2A). These findings indicate that the width of the stable microtubule overlap established by PRC1 and Kif4A is approximately equal to the sum of the end-tag lengths on both microtubules. At the lowest concentration of PRC1 tested (0.2 nM PRC1 + 6 nM Kif4A-GFP), the final overlap length was shorter than the *L*_*ET1*_ + *L*_*ET2*_, as indicated by a ratio of < 1 (Fig. 4B, black). A possible reason is that under these conditions, the end-tagged regions of the microtubules have a greater fraction of unoccupied sites that allows for further sliding and reduction in the overlap length after the collision of end-tags.

**Fig. 4.**
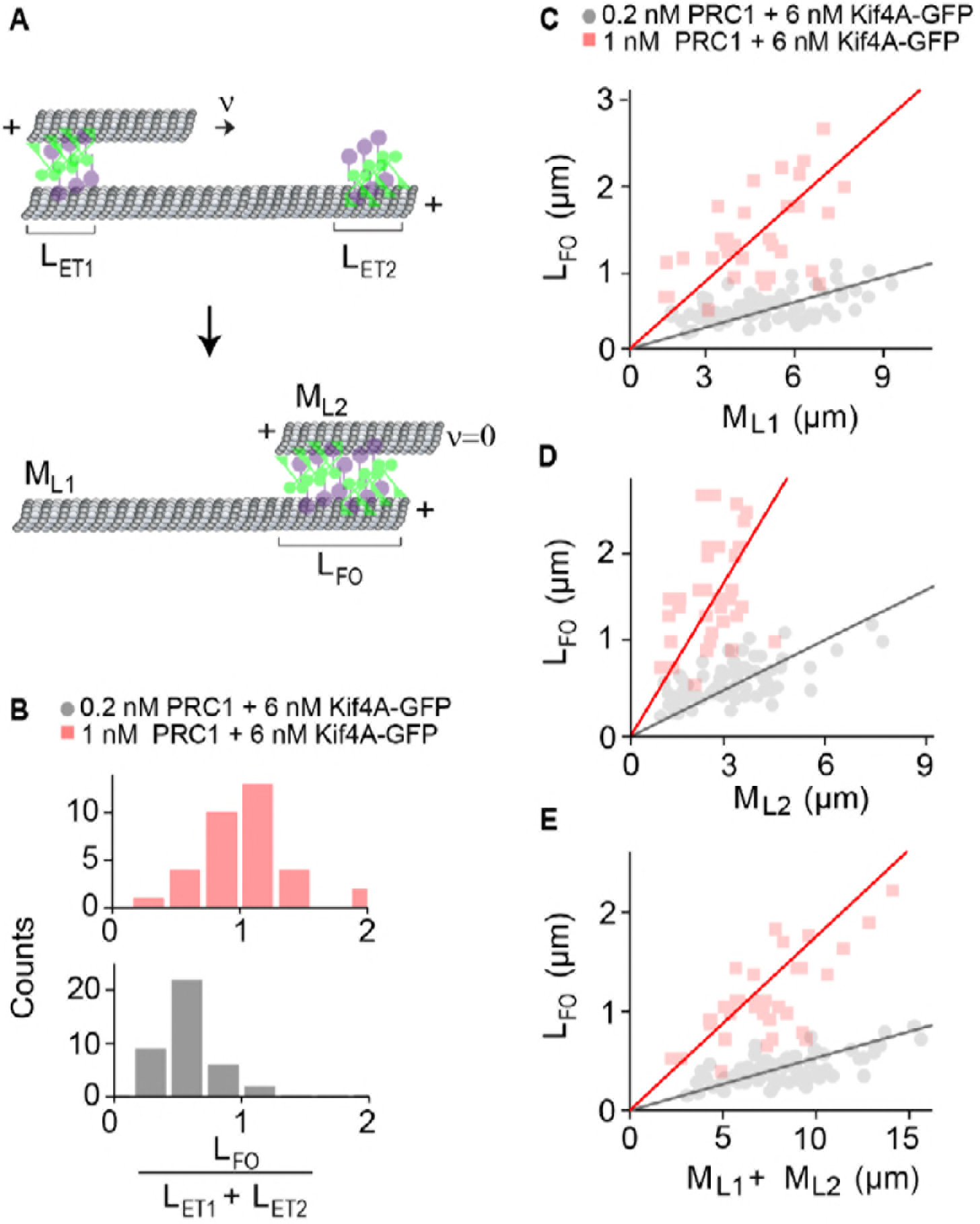
The width of the final antiparallel overlap established by PRC1 and Kif4A is determined by end-tag and microtubule lengths. **(A)** Schematic shows the formation of a stable antiparallel overlap upon collision of the two end-tags and the stalling of relative microtubule sliding. The initial overlap length is the overlap length of the moving MT on the immobilized MT at *t*=0. *L*_*ET1*_ and *L*_*ET2*_ are the lengths of the end-tags consisting Kif4A and PRC1 on the plus-end of each MT. The moving MT with length *M*_*L2*_ moves relative to the immobilized MT with length *M*_*L1*_, at velocity=*ν*. The collision and the stalling of the end-tags form a stable overlap, which is the final overlap length at *ν*=0. **(B)** Histograms of the ratio of sum of the end-tag lengths (*L*_*ET1*_ + *L*_*ET2*_) and final overlap length *L*_*FO*_. Assay conditions: (i) 0.2 nM PRC1 and 6 nM Kif4A-GFP (black; N=51) and (ii) 1 nM PRC1 and 6 nM Kif4A-GFP (red; N=33). **(C-E)** Plots of the final overlap length (*L*_*FO*_) versus (C) the immobilized microtubule length (D) moving microtubule length *(M_L2_),* and (E) and the sum of microtubule lengths (*M*_*L1*_ + *M*_*L1*_). Assay conditions: (i) 0.2 nM PRC1 and 6 nM Kif4A-GFP (black; N=75) and (ii) 1 nM PRC1 and 6 nM Kif4A-GFP (red; N=30). The Pearson’s correlation coefficient for (E) is (i) 0.65 and (ii) 0.62.

Prior work shows that the collective activities of PRC1 and Kif4A on single microtubules result in the formation of end-tags whose size scales with microtubule length [28]. This raises the question of whether the width of stable antiparallel overlap established by these proteins depends on the lengths of the two crosslinked microtubules. To examine this, we plotted the final overlap length (*L*_*FO*_) as a function of the immobilized microtubule length (*M*_*L1*_), moving microtubule length (*M*_*L2*_) and sum of both microtubule lengths (*M*_*L1*_ + *M*_*L2*_). In all three cases, we find that the final overlap length increases linearly with microtubule length (Figs. 4C-E). The slope of the line is higher at greater PRC1 concentration due to longer end-tags formed under these conditions (Fig. 4E; 0.2 nM PRC1 + 6 nM Kif4A-GFP, slope = 0.3; 1 nM PRC1 + 6 nM Kif4A-GFP, slope = 0.8). These data suggest that PRC1-Kif4A end-tags act as a barrier to microtubule sliding and establish a stable antiparallel overlap whose size is determined by the microtubule lengths.

### Examination of the mechanisms that ensure stability of the overlaps established by PRC1 and Kif4A

Why does the merging of PRC1-Kif4A end-tags during microtubule sliding result in the formation of a stable antiparallel overlap? It has been shown that the entropic forces induced by Ase1p molecules (*S. pombe* PRC1 homolog) can counter the microtubule sliding-associated forces generated by Ncd (kinesin-14) molecules to establish a stable antiparallel overlap [35]. We therefore examined if similar entropic forces are generated in the stalled microtubule overlaps established by PRC1 and Kif4A in our experiments. First, we induced the formation of stable overlaps through microtubule sliding and stalling in the presence of GFP-PRC1, Kif4A and ATP. Next, we washed the assay chamber twice with buffer containing no ATP to remove any unbound protein and nucleotide. Under this ‘no-nucleotide’ condition, we expect that the PRC1-Kif4A complexes in the microtubule overlap would essentially function as passive crosslinkers. Dual-wavelength time-lapse images were acquired for 10 mins immediately following buffer exchange. Image analysis revealed that while PRC1 was retained in the region of the microtubule overlap under these conditions, the width of the antiparallel overlap did not change during the course of the experiment (Supp. Figs. 3A-B). The lack of overlap expansion in the PRC1-Kif4A system may be due to the tight binding of the kinesin motor domain to microtubules in the absence of a nucleotide. To address this, we performed the experiment as discussed above, except the final buffer was supplemented with 2 mM ADP, a nucleotide that lowers the kinesin-microtubule affinity. As shown in Supp. Figs. 3C-F, no overlap expansion was observed under these conditions. The inclusion of 1 nM PRC1 in addition to 2 mM ADP in the final buffer also did not promote overlap expansion (Supp. Figs. 3G-H). Therefore, neither motor deactivation with ADP nor increasing the number of PRC1 molecules is sufficient to induce entropic expansions of measurable magnitude in this system, suggesting that an alternative mechanism is likely responsible for countering the Kif4A-mediated sliding forces in the antiparallel overlap.

We have previously shown that PRC1-Kif4A end-tags on single microtubules hinder motor-protein stepping [28]. Therefore, we considered if the collision of end-tags on sliding microtubules generated a stable antiparallel overlap simply by providing a steric block to sliding. To test this hypothesis, we generated stable antiparallel overlaps with PRC1, Kif4A-GFP and ATP, and subsequently exchanged the nucleotide to ADP by buffer exchange (Figs. 5A-C). As expected, no change in the overlap length was observed upon nucleotide exchange from ATP to ADP. We reasoned that under these experimental conditions, the gradual dissociation of proteins at a slow rate from the overlap should liberate a small fraction of kinesin and PRC1 binding sites on the microtubule (note: intensity analysis suggests a maximum 10% reduction of Kif4A-GFP in 2 mins). Therefore, if the moving microtubule had initially stalled due to protofilament crowding, then re-introducing ATP should allow motor-protein stepping and reinitiate microtubule sliding. To test this experimentally, we introduced buffer containing 1 mM ATP (no additional protein) into the chamber 15 mins after the ADP exchange step (Fig. 5D). We find that relative microtubule sliding is reinitiated under these conditions. Analysis of the GFP fluorescence-intensity profile at different time-points post buffer exchange revealed that new end-tags are established during microtubule sliding, which subsequently collide to establish a new stable antiparallel overlap of shorter width (Fig. 5E).

Together, these findings are consistent with a mechanism in which PRC1-Kif4A end-tags establish stable overlaps by sterically hindering the relative sliding of antiparallel microtubules. Such a ‘molecular road-block’ based mechanism also provides a simple explanation for the observed correlation between the sum of end-tag lengths and the final overlap length in this system.

**Fig. 5.**
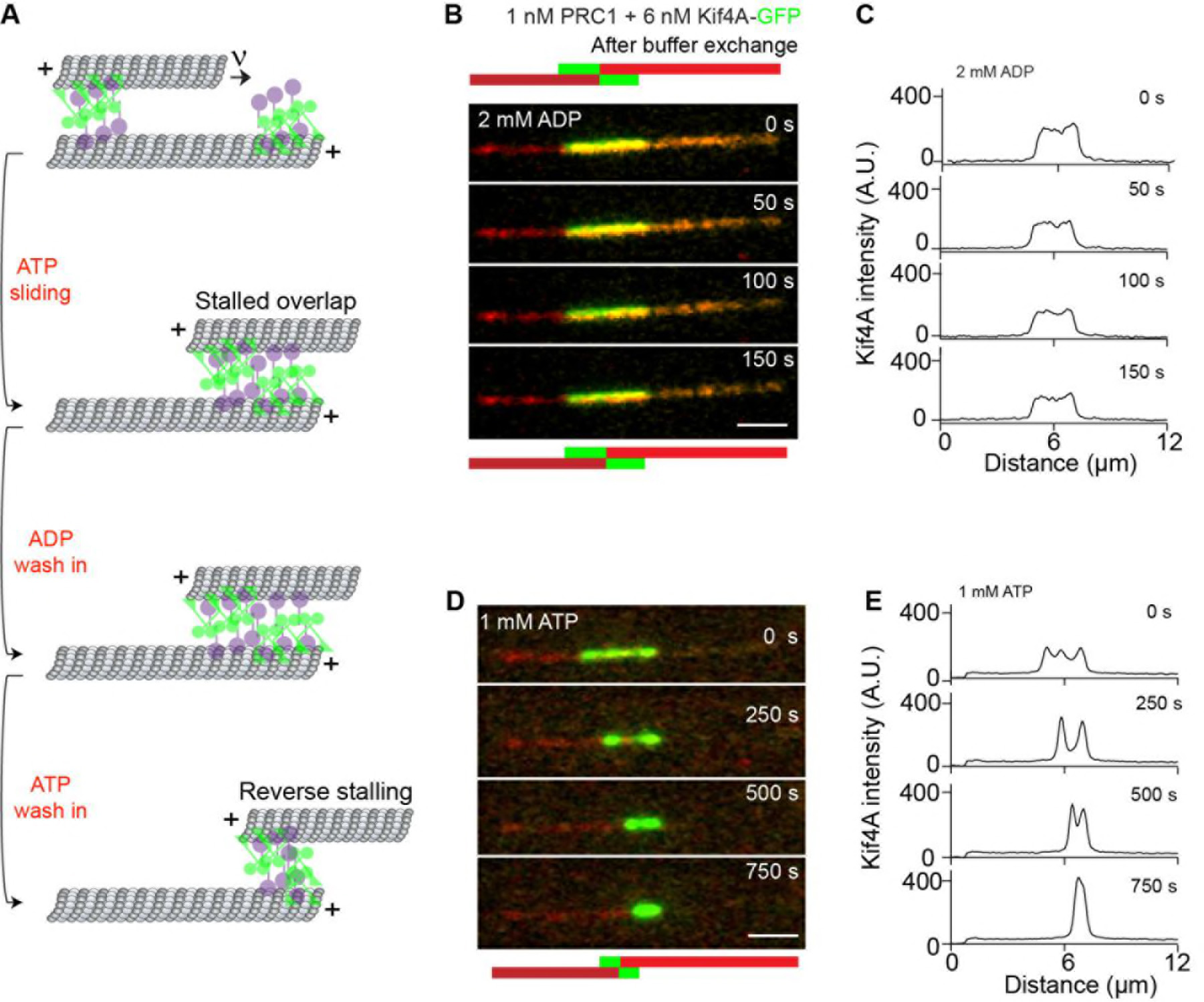
Examination of the mechanisms that ensure stability of the overlaps established by PRC1 and Kif4A. **(A)** Schematic of the ADP and ATP wash-in experiments performed with stalled microtubule overlaps (Figs. 5B-E). **(B-E)** The following figures are representative dual-channel fluorescence micrographs showing microtubules (red) and associated GFP-PRC1 (green) under different experimental conditions. **(B-C)** Time-lapse images (B) and corresponding line-scan profiles (C) of Kif4A-GFP fluorescence of a microtubule pair established as in (B) and subsequent exchange into a buffer containing 2 mM ADP. **(D-E)** Time-lapse images (D) and corresponding line-scan profiles (E) of Kif4A-GFP fluorescence of the microtubule pair in (B) after flowing in 1 mM ATP into the chamber.

### PRC1 and Kif4A align the overlap region between multiple antiparallel microtubules

How does microtubule sliding and stalling by PRC1 and Kif4A shape larger microtubule arrays? To gain insights into this question, we carefully examined the few events (N < 10) where we could clearly observe two microtubules slide relative to a single immobilized microtubule. In these events (Figs. 6A-C; 0.2 nM PRC1 + 6 nM Kif4A-GFP), we observed that both the sliding microtubules stall proximal to the plus end-tag on the immobilized microtubule. Another example of such an event in experiments with GFP-labeled PRC1 and unlabeled Kif4A is shown in Figs. 6D-F (1 nM GFP-PRC1 + 6 nM Kif4A). The data suggest that the formation of end-tags on single microtubules can establish an antiparallel array composed of multiple microtubules with closely aligned plus-ends.

We analyzed five reorganization events where we could reliably measure microtubule and overlap lengths to determine if longer microtubules result in larger final overlaps in these more complex bundles. While we cannot assess the threedimensional arrangement of the microtubules in the bundles, a simple analysis of microtubule and overlap lengths suggests that in general bundles with longer microtubules are likely to yield longer final overlaps (Supp. Fig. 4).

Together, these observations suggest that microtubule sliding and stalling by PRC1 and Kif4A can align multiple antiparallel filaments such that the region of overlap is restricted to the plus-ends of all the microtubules.

## Discussion

Pairs of crosslinked antiparallel microtubules are fundamental structural units in diverse microtubule-based architectures [1, 2]. Our findings provide insights into how the geometrical features of antiparallel microtubule arrays can be ‘decoded’ by PRC1-Kif4A complexes to govern the dynamics, stability and architecture of microtubule networks.

On the basis of our observations, we propose a mechanism for the organization of stable microtubule length-dependent antiparallel overlaps by the collective activities of PRC1 and Kif4A. PRC1 specifically crosslinks and preferentially localizes to the region of overlap between two antiparallel microtubules (Fig. 7) [25, 34]. Kif4A is recruited to the antiparallel overlap through direct interaction with PRC1 [25, 28]. The highly processive movement of PRC1-Kif4A complexes on microtubules and the slow dissociation of these proteins from microtubule plus-ends result in the formation of ‘end-tags’ on both microtubules (Fig. 7) [28]. In addition, the movement of PRC1-Kif4A complexes within antiparallel overlaps results in robust relative microtubule sliding (Fig. 7). Our results suggest that microtubule sliding occurs primarily through stepping of the motor protein in the untagged region of the microtubule overlap. During relative sliding, as the moving microtubule moves past the length of the immobilized microtubule, the distance between the end-tags at the plus-ends of both microtubules begins to shrink (Fig. 7). Microtubule movement stalls when the two end-tags arrive at close proximity during relative sliding (Fig. 7). This is likely due to the high occupancy of tubulin at end-tags, which provide steric hindrance to motor protein stepping and impede microtubule sliding [28]. An important consequence of such a ‘road-block’ mechanism is that the final overlap width is determined by the size of the end-tags formed on individual microtubules, which in turn scales with filament lengths and protein concentrations (Fig. 7).

Non-motor crosslinking proteins are primarily thought to contribute to the size and stability of microtubule arrays by opposing the active forces generated by motor proteins [39]. Microtubule organization in the PRC1-Kif4A system reveals an alternative mechanism in which a motor and a non-motor protein act synergistically on antiparallel microtubules to first promote relative sliding and then stall microtubule movement by forming a molecular roadblock. Some of the distinct features of this mechanism are as follows. First, in this system, stable arrays can be established under conditions where the non-motor:motor protein ratio may not be not sufficiently high to achieve force-balance. This is particularly advantageous in the case of interacting proteins, such as PRC1 and Kif4A, where increasing the concentration of PRC1 leads to a concomitant increase in both the levels of motor and non-motor proteins further shifting the force-balance point. Second, this system allows for robust relative sliding until the end-tags collide. This in turn leads to the establishment of stable antiparallel overlaps that are spatially restricted to microtubule plus-ends. Third, it provides a simple mechanism by which the formation of length-dependent PRC1-Kif4A end-tags on single microtubules can be readily translated to the organization of microtubule overlaps whose size scales with microtubule length.

Surprisingly, we find that the sliding velocity in the PRC1-Kif4A system is proportional to the initial antiparallel microtubule overlap length (Fig. 7). While the scaling of movement velocity with motor number has been proposed for microtubule-based transport of cargoes in the cellular cytoplasm [40–42], it is not typically observed in microtubule sliding by an ensemble of processive kinesins in *in vitro* assays. One proposed reason is the low viscous drag experienced by the moving microtubule in aqueous buffers relative to the intracellular environment, and the high magnitude of forces generated by kinesin molecules [43, 44]. Our observations in the PRC1-Kif4A system suggest that the intrinsic activity of microtubule crosslinking proteins can result in the scaling of relative sliding velocity with microtubule-overlap length even in the absence of substantial external viscous drag forces.

How might the microtubule sliding velocity scale with initial overlap length? Hints to a possible mechanism come from recent studies that investigate how the physical properties of cargoes impact microtubule sliding by motor proteins [26, 45, 46]. For example, it is observed that when kinesin motors are anchored to a diffusive lipid surface instead of a rigid glass coverslip, the gliding velocity of attached microtubules is dependent on the number of motor molecules [26]. To understand the observed scaling of velocity in the PRC1-Kif4A system, we consider the nature of the ‘cargo’ borne by the Kif4A molecule (Appendix Fig. 1). The C-terminus non-motor domain of Kif4A binds the N-terminus of dimeric PRC1 [28]. The spectrin domains at the C-terminus of PRC1 diffusively binds and crosslinks both microtubules (Appenddix. Fig. 1) [34]. Therefore the ‘moving’ microtubule is likely to be loosely coupled to the kinesin via a diffusive PRC1-microtubule linkage. In this scenario, the overall stepping efficiency is not 100%, as every 8 nm step of the kinesin does not translate into an 8 nm movement of the microtubule due to slippage arising from PRC1 diffusion. Other factors such as force dependent dissociation of PRC1-Kif4A during microtubule movement could contribute to a further reduction in the coupling between the two microtubules. We adapted the formulation developed by Grover et. al. to antiparallel sliding by PRC1 and Kif4A, and find that it can qualitatively recapitulate the velocity trend observed in our experiments (Appendix Text and Appendix Fig. 1) [46]. The modeling reveals that the diffusion constant of PRC1 and the total number of binding sites available for PRC1-Kif4A crosslinking are likely to be key parameters in determining the extent of velocity scaling with the initial overlap length (Appendix Fig. 1). Together, these analyses suggest a potential mechanism by which the velocity of microtubule sliding by PRC1 and Kif4A can scale with antiparallel overlap length.

A defining architectural feature of the spindle midzone is a stable antiparallel microtubule array with overlapping plus-ends [6, 47]. Cell-biological analyses in different model organisms indicate that both microtubule sliding and accumulation of proteins at microtubule plus ends occur on midzone arrays. The relative sliding of PRC1-crosslinked microtubules by motor proteins such as Cin8 and Kip3p in budding yeast and KLP61F in drosophila is thought to mediate the spindle elongation during early anaphase and contribute to defining the overlap width [48–50]. The accumulation of proteins at microtubule plus-ends, including multiple mitotic kinesins, is thought to effectively concentrate cytokinesis factors proximal to the site of cell cleavage [36, 47, 51]. Our biophysical analyses suggest that the geometrical features of the overlapping microtubules in the spindle may in turn tune the activity of associated proteins, and regulate the geometry and stability of the spindle midzone.

In summary, our studies show how two microtubule-associated proteins, each with its own distinct filament binding properties, can act collectively to ‘measure’ the geometrical features of microtubules arrays and ‘translate’ them to generate well-defined mechanical and structural outputs. Filament crosslinking, relative-sliding and molecular crowding are likely to represent general features of a number of biological polymers, such as actin filaments and nucleic acids, that are dynamically organized during different cellular processes. The mechanism revealed here can therefore represent general principles that regulate the size and dynamics of cellular architectures built from different polymers.

## Materials and Methods

### Protein purification

Recombinant proteins used in this study (PRC1, PRC1-GFP, Kif4A and Kif4A-GFP) were expressed and purified as described previously [28, 34].

### Microtubule polymerization

GMPCPP polymerized and taxol stabilized rhodamine-labeled microtubules were prepared with and without biotin tubulin as described previously [28, 34]. Briefly, GMPCPP seeds were prepared from a mixture of unlabeled bovine tubulin, X-rhodamine-tubulin and biotin tubulin, which were diluted in BRB80 buffer (80 mM PIPES pH 6.8, 1.5 mM MgCl_2_, 0.5 mM EGTA, pH 6.8) and mixed together by tapping gently. The tube was transferred to a 37°C heating block and covered with foil to reduce light exposure. Non-biotinylated microtubules and biotinylated microtubules were incubated for 20 mins and 1 h 45 mins, respectively. Afterwards, 100 μL of warm BRB80 buffer was added to the microtubules and spun at 75000 rpm, 10 mins, and 30°C to remove free unpolymerized tubulin. Following the centrifugation step, the supernatant was discarded and the pellet was washed by round of centrifugation with 100 μL BRB80 supplemented with 20 μM taxol. The pellet was resuspended in 16 μL of BRB80 containing 20 μM taxol and stored at room temperature covered in foil.

### *In vitro* fluorescence microscopy assay

The microscope slides (Gold Seal Cover Glass, 24×60 mm, thickness No.1.5) and coverslips (Gold Seal Cover Glass, 18×18 mm, thickness No.1.5) are cleaned and functionalized with biotinylated PEG and nonbiotinylated PEG, respectively, to prevent nonspecific surface sticking, according to standard protocols [28, 34]. Flow chambers were built by applying two strips of double-sided tape to a slide and applying to the coverslip. Sample chamber volumes were approximately 6-8 μL.

Experiments were performed as described previously [28, 34]. To make antiparallel microtubule bundles, biotinylated microtubules (referred to as ‘immobilized microtubules’ in text), labeled with rhodamine, were immobilized in a flow chamber by first coating the surface with neutravidin (0.2 mg/ml). Next, 0.2 nM un-labeled PRC1 in BRB80 + 5 % sucrose was flushed into the flow chamber. Finally, non-biotinylated (referred to as moving microtubules) were flushed in the flow cell and incubated for 10-15 mins to allow antiparallel overlap formation with the PRC1-decorated immobilized microtubules. To visualize microtubule sliding, PRC1 and Kif4A-GFP and 1 mM ATP were flowed into the chamber in assay buffer (BRB80 buffer supplemented with 1 mM TCEP, 0.2 mg/ml k-casein, 20 μM taxol, 40 mg/ml glucose oxidase, 35 mg/ml glucose catalase, 0.5% b-mercaptoethanol, 5% sucrose and 1 mM ATP) and a time-lapse sequence of images was immediately acquired at a rate of 3 frames/s. Data were collected for 10-15 mins. Key experiments and analysis were also performed with GFP-PRC1 and non-fluorescent Kif4A to rule out the effect of GFP on microtubule sliding and stalling by PRC1 and Kif4A.

All experiments were performed on Nikon Ti-E inverted microscope with a Ti-ND6-PFS perfect focus system equipped with a APO TIRF 100x oil/1.49 DIC objective (Nikon). The microscope was outfitted with a Nikon-encoded x-y motorized stage and a piezo z-stage, an sCMOS camera (Andor Zyla 4.2), and two-color TIRF imaging optics (lasers: 488 nm and 561 nm; Filters: Dual Band 488/561 TIRF exciter). Rhodamine-labeled microtubules and GFP-labeled proteins (either PRC1 or Kif4A) in microtubule sliding assays were visualized sequentially by switching between FITC and TRITC channels.

### Image analysis

ImageJ (NIH) was used to process the image files. Briefly, raw time-lapse images were converted to tiff files. A rolling ball radius background subtraction of 50 pixels was applied to distinguish the features in the images more clearly. From these images, individual microtubules sliding events were picked and converted to kymographs by the MultipleOverlay and MultipleKymograph plug-ins (J. Reitdorf and A. Seitz; https://www.embl.de/eamnet/html/bodykymograph.html). The following criteria were used to exclude events from the analysis: 1) Only kymographs where we could confidently identify exactly two microtubules in the bundle were examined further (except for the data in Fig. 6); 2) Sliding microtubules that encounter another bundle were excluded; 3) Pairs of microtubules with proximal plus-ends at initial time points could not be analyzed due to the very short duration of sliding; 4) For the sliding velocity versus initial overlap analysis, we only included kymographs where the initial overlap and the moving end-tag edge could be clearly distinguished (Figs. 2 and 3. Supp. Fig. 1); 5) For the microtubule length versus final overlap analysis, we picked kymographs both the immobilized and the moving microtubule edges could be distinguished (Fig. 4); and 6) In Figs. 3A-D, we excluded data for initial overlaps greater than 5 μm because of the existence of a few data points.

**Fig. 6.**
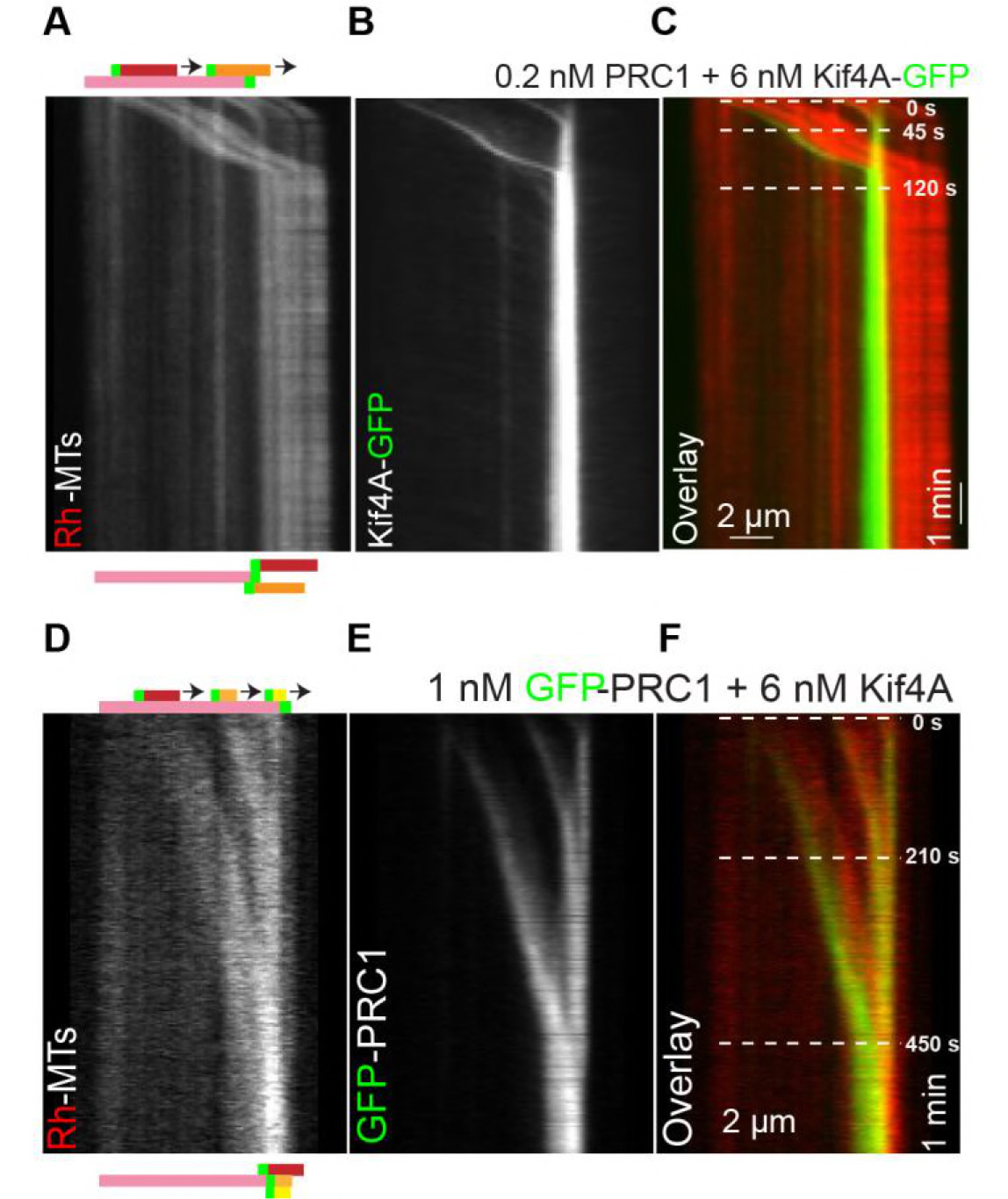
Antiparallel array composed of multiple microtubules are aligned at microtubule plus-ends formed by PRC1 and Kif4A. **(A-C)** Kymographs show the relative sliding of two microtubules relative to an immobilized microtubule (A), associated Kif4A-GFP (B) and the overlay image (red, microtubules; green, Kif4A-GFP) (C). Both moving microtubules stall at the plus-end of the immobilized microtubule. Assay condition: 0.2 nM PRC1 and 6 nM Kif4A-GFP. Scale bar: *x*: 2 μm and *y*: 1 min. **(D-F)** Kymographs show the relative sliding of three microtubules relative to an immobilized microtubule (D), associated GFP-PRC1 (E) and the overlay image (red, microtubules; green, GFP-PRC1 (F). All three moving microtubules stall at the plus-end of the immobilized microtubule. Assay condition: 1 nM GFP-PRC1 and 6 nM Kif4A. Scale bar: *x*: 2 μm and *y*: 1 min.

**Fig. 7.**
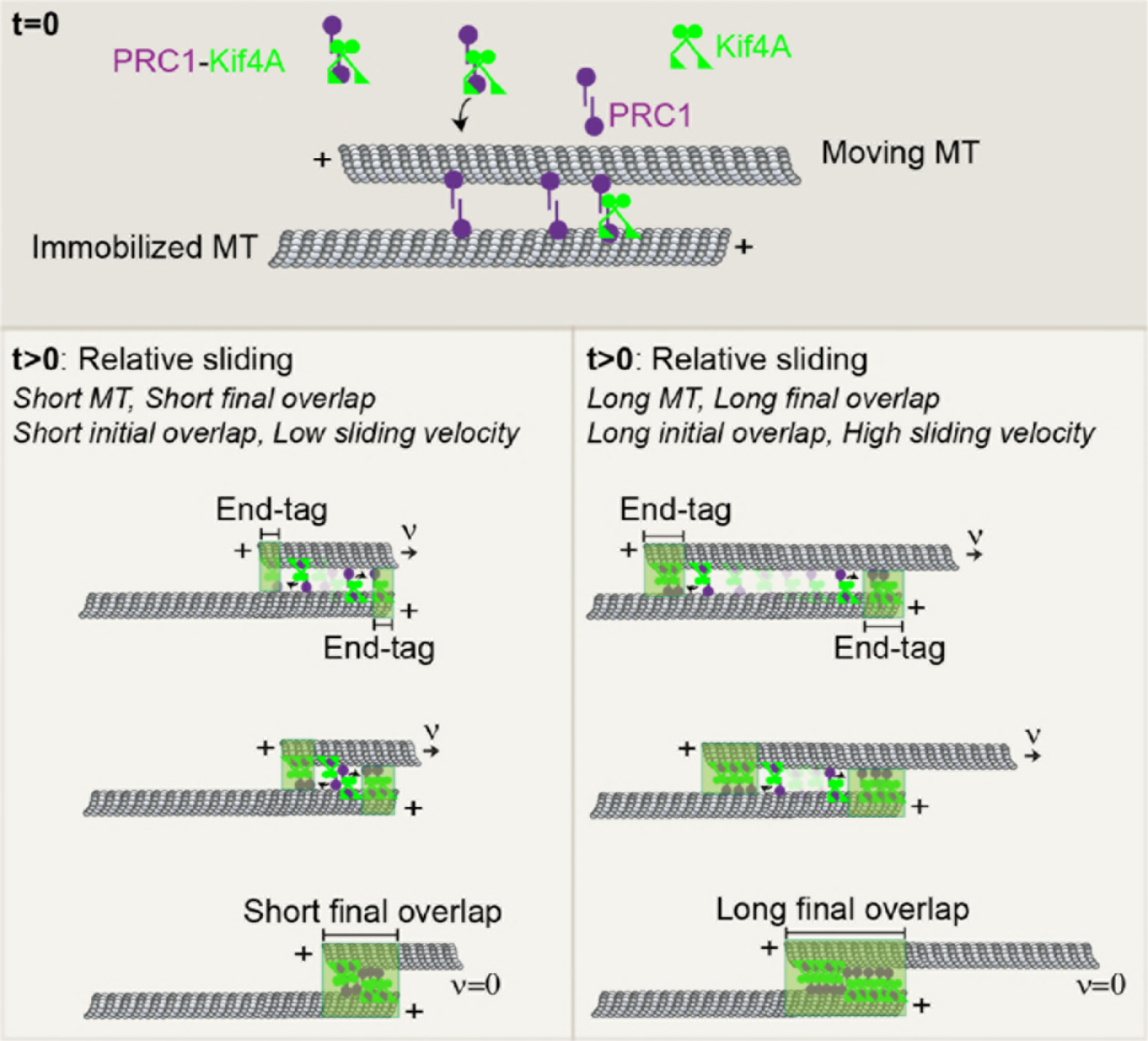
Model for the length-dependent sliding by the collective activity of PRC1 and Kif4A. Schematic shows a simple model for initial microtubule sliding and subsequent stalling of overlapping antiparallel microtubules established by PRC1 and Kif4A when the end-tags arrive at close proximity. At the initial state, *t* = 0, the ‘immobilized’ and ‘moving’ microtubules are crosslinked by PRC1 to form an antiparallel overlap. At *t* > 0, Kif4A molecules are introduced into the solution, which form a complex with PRC1. This initiates the formation of PRC1-Kif4A end-tags at the plus-ends of both microtubules as well as relative sliding of the moving microtubule. Microtubules that form shorter initial overlaps slide with lower velocity than microtubule pairs that form longer initial overlaps. A stable overlap is established when end-tags arrive at close proximity. Since the size of PRC-Kif4A end-tags scale with microtubule length, shorter microtubules form a short overlap and the longer microtubules form a longer overlap.

These kymographs were then further analyzed using a custom MATLAB program. The program first reads the input image and converts it to an array of intensity values. Next, using the ‘bwboundaries’ function, the high-intensity edges of the GFP channel kymograph were detected. If the features of the kymograph were clear, the edges of the immobilized end-tag and the moving end-tag were detected. Finally, any repeating elements due to a large amount of noise and poor contrast were removed by using the ‘unique’ function. The lines were then converted to *x*, *y* coordinates at each time point. For unclear MT or GFP channel kymographs, the kymographs can be further processed by ImageJ using the ‘Find Connected Regions’ plug-in (M. Longair; http://imagej.net/FindConnectedRegions) to distinguish the features in the kymographs more clearly. This function separates regions in the kymograph based on criteria such as having the same intensity value for the detection of edges. Afterwards, these processed kymographs can be read through the MATLAB program as described above.

To calculate the sliding velocity, the derivative of the position versus time coordinates of the external edge of the moving microtubule end-tag from the GFP channel kymograph was taken. The overlap length (*L*_*overlap*_) was measured from the MT channel. The final overlap length (*L*_*FO*_) was measured from the MT channel when the end-tags have collided and reached a steady-state. The sum of the end-tag lengths, *L*_*ET1*_ and *L*_*ET2*_, were determined by measuring the end-tag length before the collision of the end-tags from the GFP channel. The sum of the microtubule lengths, *M*_*L1*_ and *M*_*L2*_, were measured from the rhodamine channel, and their sum was also plotted.

**Data availability:** The data that support the findings of this study are available from the corresponding author upon request.

**Supplementary Information:** The supplementary information contains Supp. Figs. 1-4. Appendix: The appendix contains the theoretical description of the scaling of sliding velocity with initial overlap and appendix references.

## Acknowledgements

The authors would like to thank Tarun Kapoor (Rockefeller Univ.) for generous support during initial stages of this project. The authors would also like to thank Scott Forth (RPI), Doug Martin (Lawrence College) and Tarun Kapoor (Rockefeller Univ.) for helpful comments.

### Author Contributions

R.S. conceived and designed the project. S.S.W and R.S. performed experiments, analyzed the data and wrote the manuscript.

### Competing financial interests

The authors declare no competing financial interests.

## Appendix 1. Theoretical description of the scaling of microtubule sliding velocity with initial overlap length

We propose a simple model consistent with our experimental observations that the microtubule sliding velocity in the PRC-Kif4A system depends on the initial antiparallel overlap length (Figs. 3A-B). The central feature of this model is that sliding is mediated by PRC1-Kif4A molecules that can diffuse along one microtubule and step along the other. As shown in the schematic in Appendix Fig. 1A, the non-motor crosslinker PRC1 (purple) diffuses one-dimensionally along the microtubule and the motor kinesin molecule (green) steps along the lattice to the plus-end of the microtubule. This will result in a reduction in the coupling between kinesin stepping and translocation of the moving microtubule. Therefore, the net sliding velocity is lower than the velocity of Kif4A stepping on single microtubules (Appendix Fig. 1A). This configuration is conceptually similar to the sliding of microtubules by lipid membrane-anchored kinesins [1]. Therefore we adapt the theoretical framework developed by *Grover et al.* to examine the dependence of microtubule sliding velocity by PRC1-Kif4A complexes on initial overlap length of the array [1].

For this system, we define: the stepping velocity of the motor as *ν*_*step*_ which propels the microtubule with *ν*_*MT*_ and *ν*_*slip*_ as the net ‘slipping rate’ of the PRC1-Kif4A on the microtubule due to PRC1 diffusion and other potential factors such as force dependent dissociation of PRC1 and Kif4A, thus

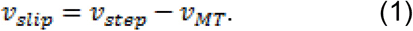

Next, for simplicity, it is assumed that the microtubule-PRC1-Kif4A complex is at equilibrium force. Therefore, the net force acting on the system is zero. The force balance equation becomes

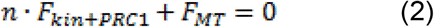

where *n* is the number of molecules in the overlap, *F*_*kin*+*PRC1*_ is the net frictional force acting on the kinesin due to diffusion of PRC1 molecules on microtubules, and *F*_*MT*_, is the external force acting on the microtubule due to hydrodynamic drag of the solution.

The frictional force on a PRC1 molecule that is dragged by a kinesin motor with velocity is *F*_*kin*+*PRC1*_ = γ*ν*_*slip*_, where 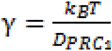 where *D*_*PRC1*_= 0.29×10^4^ nm^2^/s for PRC1 molecules in a densely-decorated microtubule overlap [1].

The frictional force on a moving microtubule, *F*_*MT*_ = γ_MT_*ν*_*MT*_ where *ν*_*MT*_ is the velocity of the microtubule, 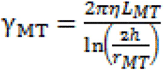 is the drag coefficient for a cylindrical object moving parallel to the surface, ***η*** is the viscosity of water, *L*_*MT*_ is the length of the microtubule, *h* is the height of the microtubule, and *T*_*MT*_ is the radius of the microtubule.

Because microtubule has 13 protofilaments, the molecules between two microtubules can theoretically bind to more than one protofilament. For this reason, the maximum number of molecules *n* in an overlap *l* can be written as 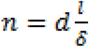 where *d* is the number of protofilaments and the length of a single site, *δ* = 8 nm. If the fractional occupancy is *a*, then we can rewrite Eq. 2 as:

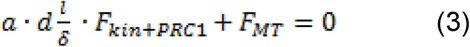

where *a* is the fractional occupancy.

Plugging in the *F*_*MT*_ and *F*_*kin*+*PRC1*_, Eq. 3 becomes,

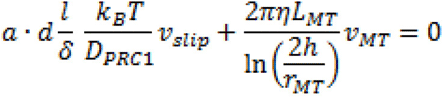

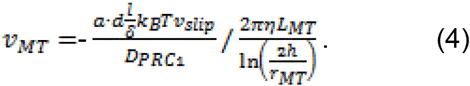

Substituting equation (1) in (4), we yield

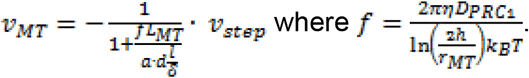

Using previously published parameters (Table A1), we examined which factors are likely to dictate the dependence of initial overlap length on sliding velocity. We find that the theoretical model predicts that the sliding velocity as a function of overlap depends weakly on the moving microtubule length (Appendix Fig. 1B) and the (ii) fractional occupancy of the molecules (Appendix Fig. 1C), but depends strongly on the: (i) the diffusion constant (Appendix Fig. 1D) and (ii) the number of protofilaments that are available for cross-bridging (Appendix Fig. 1E). Consistent with these trends, in our analysis of the experimental data (Fig. 3), we observed a stronger dependence of sliding velocity on initial overlap length at higher Kif4A-PRC1 ratios where we would expect a greater percent of complexes in the overlap contributing to sliding.

**Table A1:**
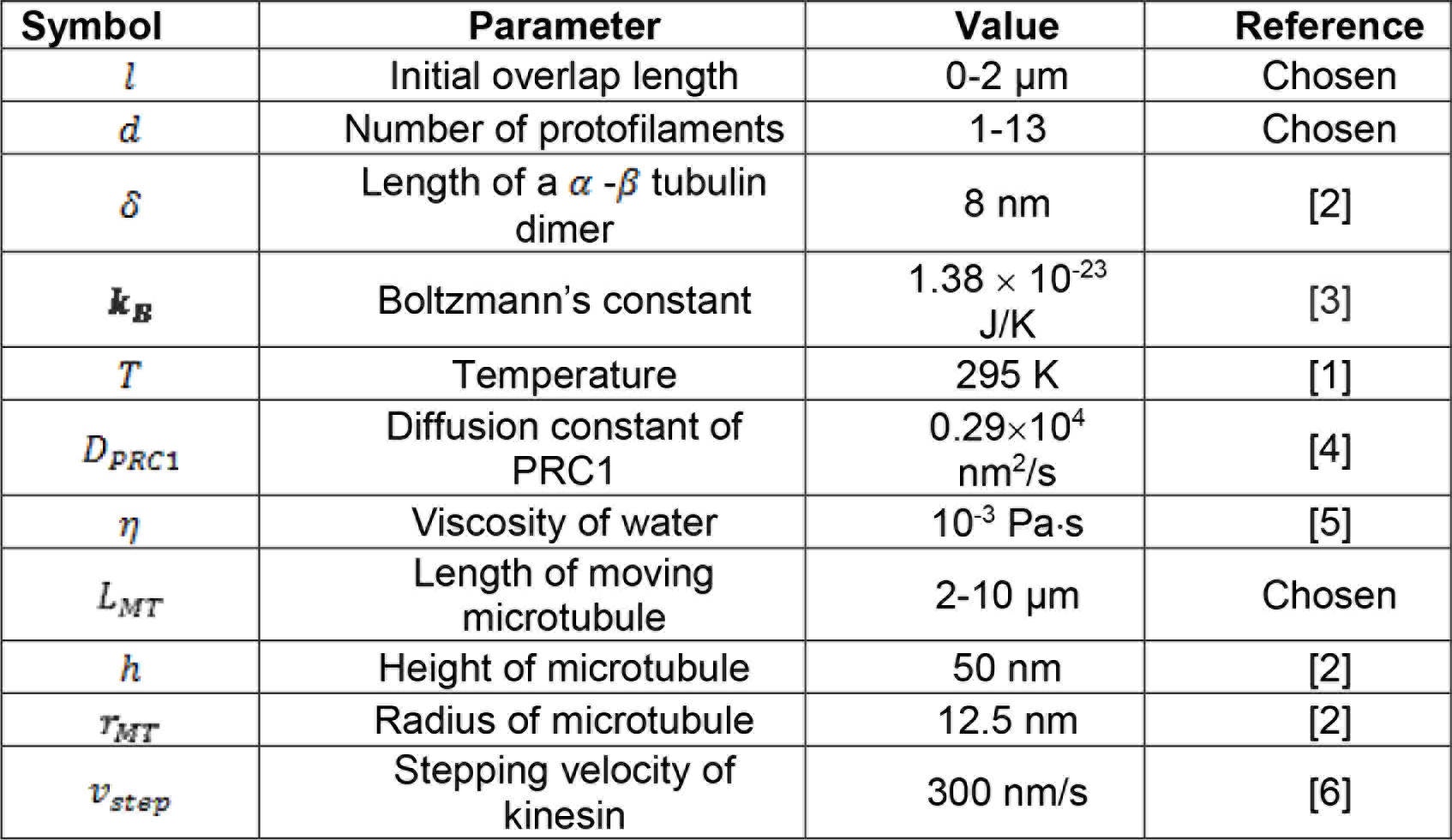
Parameter values

| Symbol | Parameter | Value | Reference |
| --- | --- | --- | --- |
| $l$ | Initial overlap length | 0-2 $\mu\text{m}$ | Chosen |
| $d$ | Number of protofilaments | 1-13 | Chosen |
| $\delta$ | Length of a α-β tubulin dimer | 8 nm | [2] |
| $k_B$ | Boltzmann's constant | $1.38 \times 10^{-23}$ J/K | [3] |
| $T$ | Temperature | 295 K | [1] |
| $D_{\text{PRC1}}$ | Diffusion constant of PRC1 | $0.29 \times 10^4$ nm<sup>2</sup>/s | [4] |
| $\eta$ | Viscosity of water | $10^{-3}$ Pa·s | [5] |
| $L_{\text{MT}}$ | Length of moving microtubule | 2-10 $\mu\text{m}$ | Chosen |
| $h$ | Height of microtubule | 50 nm | [2] |
| $r_{\text{MT}}$ | Radius of microtubule | 12.5 nm | [2] |
| $v_{\text{step}}$ | Stepping velocity of kinesin | 300 nm/s | [6] |

**Appendix Figure 1.**
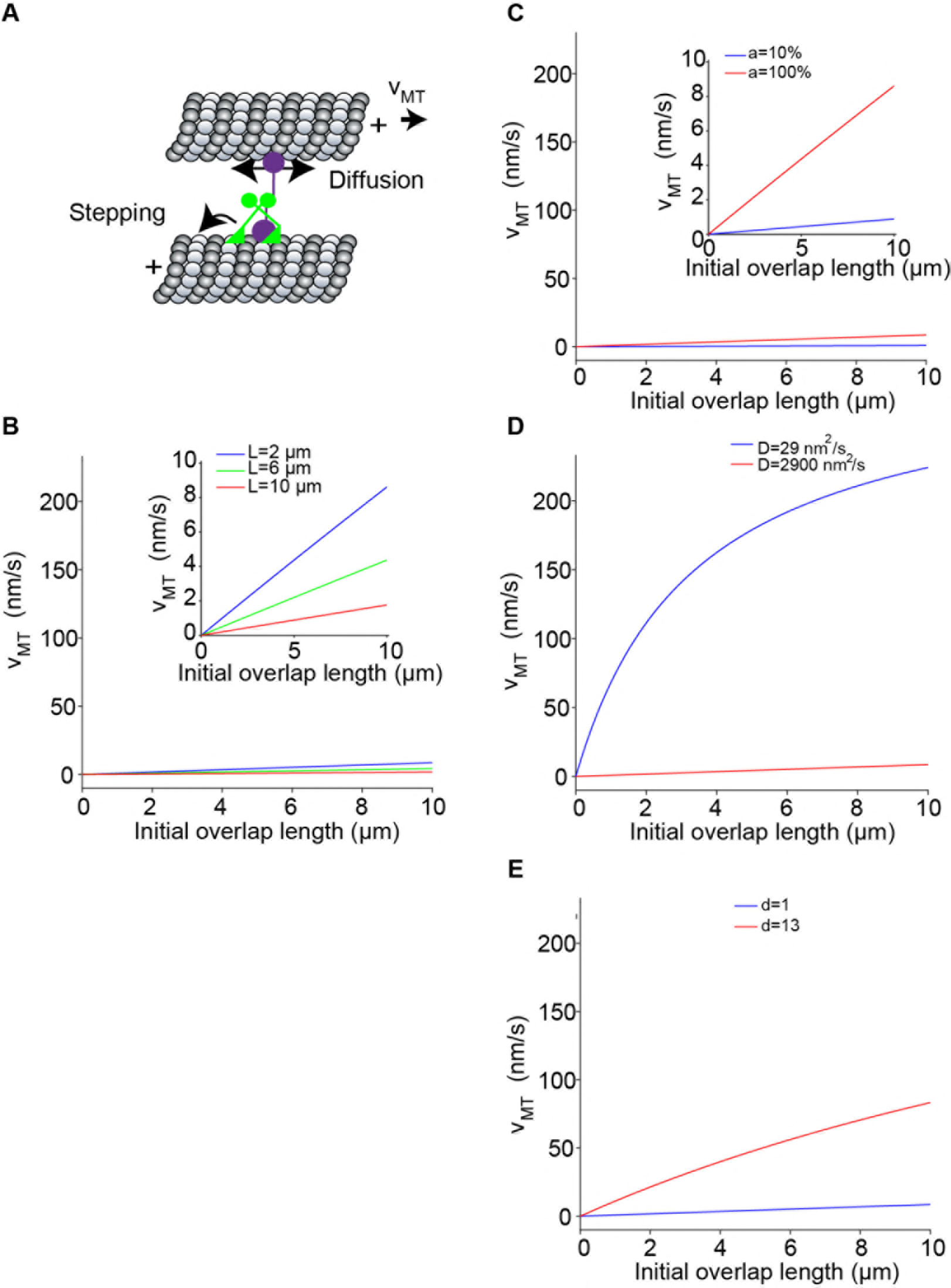
Box initially at rest on sled sliding across ice.

Theoretical model of the scaling of microtubule sliding with the initial overlap length. **(A)** Schematic shows a simple model of microtubule sliding driven by a microtubule anchored motor-non-motor complex. The non-motor PRC1 molecule (purple) diffuses one-dimensionally along the microtubule and the kinesin molecule (green) drags the PRC1 molecule and steps along the lattice, which results microtubule sliding with velocity *V*_*MT*_. **(B-E)** Plot of the sliding velocity as a function of initial overlap length color-coded by (B) moving microtubule lengths (blue: 2 μm; green: 6 μm; red: 10 μm), (C) fractional occupancy, *a* (blue: 10%; red: 100%), (D) diffusion constant, ***D***, (blue: 29 nm^2^/s; red: 2900 nm^2^/s), and (E) number of protofilaments available for crosslinking and stepping, *d*.

